# MicrobioRel: A Set of Datasets for Microbiome Relation Extraction

**DOI:** 10.1101/2025.08.03.666357

**Authors:** Oumaima El Khettari, Daniel Batteux, Solen Quiniou, Samuel Chaffron

**Author notes:** Corresponding authors: Oumaima El Khettari, Solen Quiniou and Samuel Chaffron.

## Abstract

Biomedical knowledge curation relies on a variety of Natural Language Processing tasks, including biomedical entity recognition and document-level relation extraction. With the growing size and capabilities of Language Models, effectively deploying them in specific and specialised domains remains a persistent challenge, highlighting the need for high-quality, domain-adapted datasets. In this work, we present MicrobioRel, a corpus of two datasets to study the relations between biological entities in the gut microbiome. The first dataset, MicrobioRel-cur, is a document-level labelled corpus, corresponding to paragraphs from journal articles that were manually annotated with different types of relations between biomedical concepts in the gut microbiome domain. We describe its creation process, annotation guidelines, and key statistics. On this dataset, we evaluated different architectures for relation extraction and identify PubMedBERT as the most effective model for this task. We also created a second dataset, MicrobioRel-pred, by generating relation predictions on other journal articles using the fine-tuned PubMedBERT model. We demonstrate its potential to extract meaningful interactions. The MicrobioRel is a crucial resource for advancing tasks like automatic knowledge extraction in specialised domains such as the gut microbiome, facilitating hypothesis generation and supporting scientific discovery.

## Introduction

On the one hand, knowledge bases serve as fundamental resources for clinicians and practitioners, supporting the development of therapies, diagnostic tools and scientific theories [38]. Maintaining accurate and up-to-date knowledge bases ensures that the latest advances in the biomedical field are accessible. Regular updates are thus essential to ensure that these repositories reflect the most up-to-date understanding of biological processes and disease mechanisms. On the other hand, the volume of biomedical literature has increased substantially in recent years, presenting both an opportunity and a challenge to harness this expanding scientific data. This flood of information offers the potential for knowledge discovery, but requires effective methods to extract meaningful key insights. Extracting information from scientific articles, particularly recent ones, allows, for example, the categorisation of associations between multiple concepts. This categorisation can then support the automatic completion of existing knowledge structures or biomedical ontologies [30].

As manual curation and traditional literature reviews struggle to keep pace, more efficient and automated methods are needed to extract and categorise the relations embedded in biomedical documents [38]. Large Language Models (LLMs) have the potential to revolutionise information synthesis from scientific literature, particularly for tasks such as document-level relation extraction in the biomedical domain [20]. While previous studies have emphasised sentence-level relation extraction, such as protein-protein interactions [19], this approach limits contextual understanding by overlooking inter-sentence relations. Furthermore, specific annotated datasets in the biomedical domain are scarce, complicating the task of capturing diverse semantic relations [7]. Despite recent progress, specialised corpora remain essential for effective knowledge acquisition. One area of growing interest in biomedicine is the gut microbiome, an ecosystem of diverse and interacting microorganisms that shape health and disease of the host throughout life. Understanding how this microbiome influences human health is critical [27] but no labelled corpus on this domain is available.

Several corpora have been developed for relation extraction (RE), mainly at the sentence level. These corpora, such as Protein-Protein Interactions (PPI) [19], AIMed [5], and Drug-Drug Interactions (DDI) [22], focus on specific pairs of entities. Extensions include DDI13 [42] for drug-drug interactions and ChemProt [28] for chemical-protein interactions. However, these datasets rely heavily on distant supervision [35], which can introduce label noise. In the microbiome field, MiDRED [23] provides 3,116 annotated microbe-disease relations. All of these sentence-level corpora lack document-level semantic context. While BioRED [32] includes inter-sentence relations, it excludes entity types such as species.

To perform relation extraction, early methods relied on feature-based machine learning models such as SVM [46] and Random Forest [4], which performed well with engineered linguistic features. Neural networks, including CNNs and RNNs [17, 49], improved context handling but struggled with long distance dependencies. Transformer-based models such as BERT and ATLOP [55] addressed these challenges by leveraging contextual embeddings. Generative frameworks, including seq2seq models, have also been explored for end-to-end RE [15]. Despite these advances, a comprehensive comparison of these architectures in low-resource and specialised domains, such as the microbiome, remains underexplored.

In this work, we present MicrobioRel, a publicly available set of two datasets^1^, which can be used to study the relations between biological entities in the gut microbiome. The first dataset, MicrobioRel-cur, is annotated for document-level relationship extraction, covering 6 entity types and 22 relationship types within the human microbiome domain. The corpus is based on carefully selected scientific articles related to the gut microbiome. MicrobioRel-cur allows us to perform a comparative performance analysis of relation extraction models, ranging from Support Vector Machines to Llama3, on a low-resource specialised domain that is the gut microbiome. The second dataset, MicrobioRel-pred, was created by generating relation predictions on other journal articles using the best performing relation extraction model, corresponding to a fine-tuned PubMedBERT model. In addition, to illustrate the relevance of MicrobioRel-pred, we present a case study of extracted relations related to Inflammatory Bowel Disease (IBD). From these extracted relations in MicrobioRel-pred, we built an entity-relation graph, for which we provide a detailed analysis and discussion.

## Materials and Methods

### Document Selection

The scientific articles used in this study were curated from the PubTator collection of articles hosted by the PubMed Central (PMC) repository^2^. In order to specifically target articles relevant to the gut microbiome, a selection was performed through a query composed of Medical Subject Headings (MeSH) terms [31] directly related to the topic of interest. The query was as follows *<*MESH:D005767 OR (MESH:D000069196 AND MESH:D064307) *>*. This data collection process was performed on November 2022, and resulted in 23 216 articles. Table 1 shows the MeSH identifiers along with the corresponding terms used in this study.

**Table 1.** MeSH terms and their labels

| MeSH Identifier | Corresponding label |
| --- | --- |
| D064307 | Microbiota |
| D000069196 | Gastrointestinal Microbiome |
| D005767 | Gastrointestinal Diseases |

Although the use of MeSH terms can be an effective means of selecting articles on a particular topic, it lacks precision and often results in a very large number of articles. As a solution, we defined a set of heuristics based on the identified named entities within the articles to rank them. The heuristics consisted of the total number of annotations and the number of unique annotations for each entity type. Using this information we then ranked documents based on maximising the number of Species and Disease mentions and unique mentions, as these are fundamental elements of currently under-represented interactions.

The 262 selected paragraphs for this study originate from 2 journal articles that includes an important number of annotated species and diseases. The first article focuses specifically on the role of the gut microbiota in health and cardiovascular diseases [47] and the second encompasses research on the study of connections between the gut metabolome and neurological disorders, an aspect of microbiome study that is currently being highly studied [34].

### Annotation Process

We started the annotation process using the concepts provided by PubTator [48] by selecting the following entities: Species, Disease, Chemical, Gene, Cell Line and Mutation. We used PubTator [8] as our primary resource for sourcing scientific articles, as it provides Named Entity Recognition (NER) annotations for concepts that align with our specific areas of interest. We used the Species entities pre-annotated in PubTator [14], which is particularly useful for our study of the gut microbiome. The latter is a concept that encompasses a wide range of organisms, including bacteria, fungi and viruses. Here we used the Species entities pre-annotated in PubTator [14], as they allow us to identify and track the various organisms that form the gut microbiome. This is particularly useful since the gut microbiome encompasses a wide range of species, including bacteria, fungi, and viruses. In the PubTator format, articles are divided into paragraphs, that may be single sentences, each of which representing a unit of meaning. This suggests that the relation could be articulated anywhere within the paragraph, whether in the same sentence or across sentences.

#### Entity Curation

As annotations provided by PubTator may be incorrect, the annotation process allows for the correction of annotated entities, including deletions, modifications and additions. Annotations should not be based on external knowledge. Therefore, information should be explicitly expressed in the text, whether it concerns entities or interactions.

The curation process consists of two main steps: revising the entity annotations proposed by PubTator (curation), and annotating the relations. While the annotation of named entities is not the primary focus of our work, we recognise its key role in the accuracy of subsequent tasks. In order to mitigate potential discrepancies, we have implemented a correction phase, which is based on defined concepts and rules set out in the annotation scheme. The guidelines and rules are detailed in the annotation scheme, which is included with the MicrobioRel dataset and freely available online (on GitHub).

#### Relation Annotation

The selected interaction types between pairs of entities are partly derived from the UMLS Semantic Network [3]. They describe interactions that can potentially occur between the previously mentioned entities. Our list of 22 relations is defined according to a hierarchical level and is as follows Increase, Decrease, Stop, Start, Improve, Worsen, Possible, Presence, Negative correlation, Affects, Causes, Complicates, Experiences, Interacts with, Location of, Marker/Mechanism, Prevents, Reveals, Treats, Physically related to, Part of, Associated with.

To facilitate the annotation process, a decision tree has been created and is included in the annotation scheme. It brings together the annotation guidelines and the annotators’ observations. It provides guidance depending on the type of relation and the entities involved. For example, a specific section is dedicated to entities labelled Disease, where the annotator must decide whether the described relation modifies or simply influences a disease. For each relation, examples are provided in verb format to illustrate semantic expressions. The use of the decision tree in the annotation process not only increased consistency and accuracy, but also reduced the average annotation time per paragraph by at least 10 minutes.

#### Annotation Tool

For the annotation task, we used the web-based collaborative tool Label Studio [44]. The tool allows annotators to modify or delete PubTator annotations as needed, while simultaneously annotating the relations between them, as shown in Figure 1. In this example, the entity *microbiota* has been manually annotated as a Species, as it has not been annotated by PubTator. In addition, the condition *dysbiosis* is involved in two relations, each involving a chemical (*BCAAs* and *aromatic amino acids*). This illustrates that an entity can be involved in several relations within the same paragraph or sentence.

**Fig. 1.**
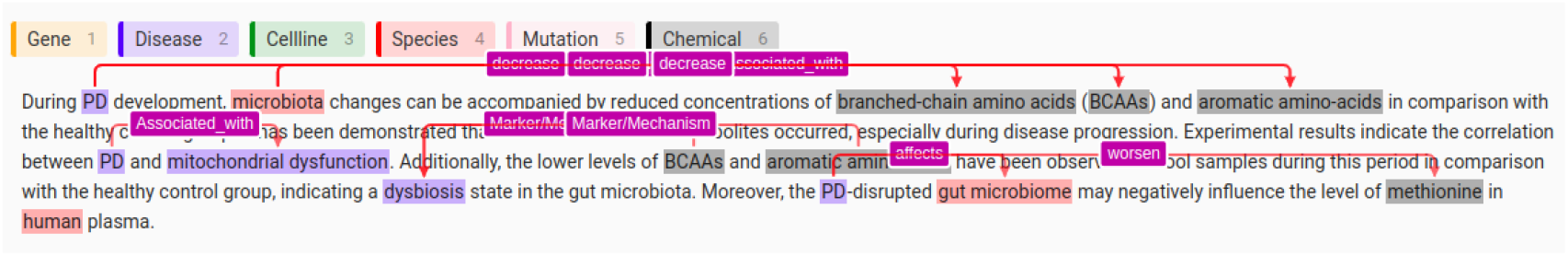
Example of a paragraph annotated with 14 entities and 9 relations in Label Studio. Notable relations include: Parkinson’s disease (PD) is associated with changes in the microbiota and mitochondrial dysfunction; Chemicals (branched chain amino acids, BCAAs) decrease during PD development; PD disrupts the gut microbiome, potentially affecting human plasma methionine levels.

### Relation Extraction Task

Here, we modelled the task of relation extraction under two paradigms: a supervised classification task and an unsupervised generative task. The goal is to define the class of the relation between known entities, using traditional machine learning models as well as transformer-based approaches to evaluate the performance under specialised low-resource settings.

#### Data Pre-Processing

The first pre-processing step consists in adding annotated examples of entity pairs with no relation between the entities. Intuitively, and before labelling a relation, it is important to check for its existence. Labelling the non-existence of a relation can be a difficult task, as the probability of having no relation between two entities is significantly higher than the probability of having one. To deal with this case, two approaches are possible. The first involves a two-step process where we first classify the presence or absence of a relation and then classify the detected present relations with their specific type. The main drawback of this approach is the high risk of error propagation from the first step to the second step [51]. Alternatively, in the second approach, a None type is added to the set of possible relation types for entity pairs without a relation. To select examples of entity pairs with a None label, each paragraph is parsed to calculate all possible entity pairs. Those with a relation label are then extracted before a selective sampling of the remaining pairs is performed, ensuring that the None label accurately represents the majority class to reflect real world scenarios. We have therefore chosen this second approach.

The second pre-processing step is to create a context for each relation instance. Since a single paragraph can contain multiple relations between different entities, we created as many duplicates of the paragraph as there are relations in it, with each duplicate containing only one of the relations.

Finally, 70 % of the dataset is set for training, and the remaining 30 % is split equally between the development set and the test set. See Section 3 for more details.

#### Supervised RE Models

As a baseline, we performed RE using widely-used statistical Machine Learning algorithms, namely Support Vector Machine [9] and Random Forest [4]. In addition, we fine-tuned selected BERT-based models, including baseline BERT [12], BioBERT [29], SciBERT [2], PubMedBERT [18] and BioLinkBERT [52], using the MicrobioRel training and validation sets. We also applied the ATLOP model [55] (an architecture designed for document-level RE), and its derivative DREEAM [33]. Both models were fine-tuned on the MicrobioRel training and validation sets. To further explore different architectures, we experimented with fine-tuning encoder-decoder models, specifically Flan-T5 [6] and BioBART [53], using the same fine-tuning technique as for the previous models.

#### Unsupervised Generative RE Models

Finally, we modelled the RE task as a generative problem by instructing the Llama2 [45] and Llama3 [13] models to generate the relation between two given entities, without any fine-tuning, and only performed the task as a generative task on the test set. We also include Mistral 7B and its biomedical version BioMistral. LLMs were used in inference mode, employing top-k sampling (top k=10), and a temperature of 0.1 for controlled randomness. For each prompt, a single output sequence was generated, with generation stopping upon reaching the end-of-sequence token.

## Results

### MicrobioRel-cur: A Curated and Annotated RE Dataset for the Gut Microbiome

#### Entity-Wise Analytics

Table 2 sums up the count of named entities after curation. Notably, in 67.93% of the paragraphs, modifications have been performed, with respect to the entities imported from PubTator. These alterations include corrections of imported annotations as well as the addition of forgotten entities. In the majority of instances, corrections have been applied to composed named entities that were only partially annotated. Additionally, adjustments were made to plural forms of entities that were not initially recognized, even after adding the ‘s’ to indicate plurality. Hence, 1 056 new named entities have been added to the corpus (representing 36.2 % of the total number of entity mentions).

**Table 2.**
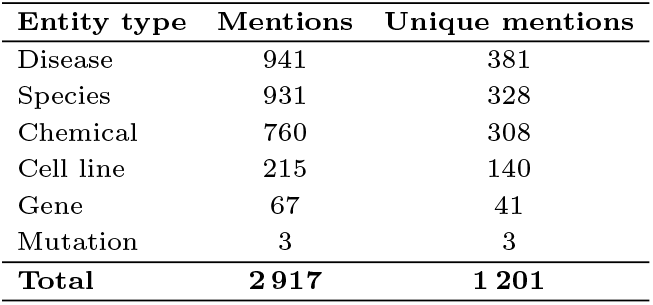
Named entity count on the corpus.

| Entity type | Mentions | Unique mentions |
| --- | --- | --- |
| Disease | 941 | 381 |
| Species | 931 | 328 |
| Chemical | 760 | 308 |
| Cell line | 215 | 140 |
| Gene | 67 | 41 |
| Mutation | 3 | 3 |
| **Total** | **2 917** | **1 201** |

#### Relation-Wise Analytics

Relations are entirely sourced from the annotation process. It resulted in 1 944 relations within the 262 paragraphs, distributed across classes as shown in Table 3. As we annotated texts in their original form, it leads to imbalanced classes.

**Table 3.** Relation label counts in the MicrobioRel corpus

| Relation type | Count | Relation type | Count |
| --- | --- | --- | --- |
| Associated_with | 210 | Possible | 56 |
| Part_of | 204 | Worsen | 51 |
| Affects | 202 | Marker-Mechanism | 41 |
| Increase | 185 | Prevents | 39 |
| Location_of | 146 | Physically_related_to | 31 |
| Causes | 144 | Negative_correlation | 23 |
| Experiences | 132 | Treats | 17 |
| Decrease | 121 | Complicates | 10 |
| Start | 118 | Presence | 9 |
| Improve | 102 | Reveals | 7 |
| Interacts_with | 86 | Stop | 6 |

It is not surprising that the most prevalent one is Associated with, commonly employed when the nature of the relation is ambiguous. We report an average of approximately 10 semantic relations per paragraph, suggesting a possible correlation between the number of relations and the paragraph length, on account for the fact that relations are semantically represented. And indeed, we observed a positive correlation between paragraph length and number of relations (Figure 2). Nevertheless, it is important to mention that the relationship between the number of words in a text and the probability of having more semantic relations between entities is not a strict rule; it can vary depending on the specific context and content of the text. In our case, academic writing style and journal articles are considered dense in information, hence the high prevalence of annotated relations.

**Fig. 2.**
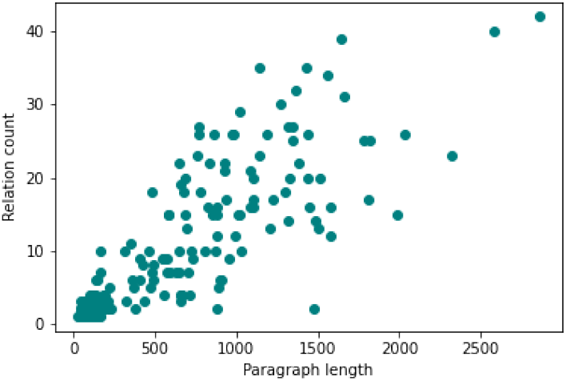
Scatter plot showing the relationship between paragraph length and the number of annotated relations in MicrobioRel. Longer paragraphs generally contain more relations, with most paragraphs clustering under 1000 characters and 10 relations, while a few outliers exhibit higher counts.

#### Inter-Annotator Agreement

In order to validate the reliability of the annotation and the accuracy of the guidelines, 32 paragraphs of the corpus have been annotated by 3 annotators. One annotator had specialized expertise in the biomedical field and prior experiences with the studies of species interactions. A second annotator had sufficient domain knowledge and received specific training for the task. The third annotator had more limited domain expertise but is well aware of the task of relation extraction. This setup allowed us to assess both the clarity of the guidelines and the influence of domain expertise on annotation quality. The standard method for evaluating inter-annotator agreement (IAA) in classification tasks are Cohen’s Kappa and Fleiss’s Kappa. Herein, we did not use these well-known metrics due to the nature of the annotations and the task. These metrics assume annotators evaluate the same set of items, whereas in this case, annotators did not annotate the same number of elements (representing relations), leading to differences in coverage. Moreover, these metrics tend to penalize missing annotations, which could unfairly lower agreement scores. Along the same lines, the work done by [26] and [11] have demonstrated that these metrics are not the best suited especially for the task of annotating relations. Consequently, studies have shown that the F-measure approximates the kappa, by considering one annotation as the reference [54].

The analysis yielded an F1-score of approximately 51%, when considering the exact match of entire triplets. When considering only the participating entities, it raised to 78%, suggesting that the annotators agree on the existence of links between entities, and have disagreements on the type of relations to choose. An analysis of the relations they disagree on reveals that their differences are primarily due to selecting relations with varying levels of precision, rather than identifying relations that are contradictory or not relevant to the context.

For the 32 paragraphs used to compute inter-annotator agreement, annotations from the most specialized annotator were retained. For the remaining 230 paragraphs, the annotations from a second annotator, who has sufficient domain expertise for the task, were used.

### Relation Extraction on MicrobioRel-cur

#### Experimental Setup

For the supervised approach, we fine-tuned the cased version of the base models of the mentioned BERT-based architectures over 30 epochs, using a fixed batch size of 4 and a learning rate set to 1 *×* 10^*−*5^, except for Flan-T5, where the small version was used. For the machine learning approaches, we used Word2Vec to encode feature representations for both the Support Vector Machine and Random Forest classifiers. Word2Vec was trained on the annotated triplets to capture vector representations of the relations and entities. In the case of SVM, given the multiclass nature of the task, a one-vs-one classification strategy was employed to handle the multiple relation categories.

For the unsupervised generative experiments in this study, a prompt was designed to classify relationships between two entities based on the input text. The prompt follows a structured format, consisting of task instructions, output constraints, and the input text. This approach, which has been shown to be effective in various studies [36, 41], is suitable for information extraction tasks. The experiments were conducted in a zero-shot setting, without examples in the prompt. This choice was based on preliminary experiments, which showed that including a limited set of relations in examples introduced bias, as the model tended to over-generate the demonstrated classes. Given the 23 possible relation classes and the token size limit, a zero-shot approach was preferred. Table **??** shows an example of the prompt.

The performance of the RE models was evaluated using weighted Precision (P), Recall (R), and F1-scores on the test set, with the results summarized in Table 4.

**Table 4.** Weighted Precision, Recall, and F1-scores (in %) of relation extraction models on the MicrobioRel test set. Results in bold are the highest scores. ^*†*^LLMs are used in zero-shot inference without fine-tuning.

| Model | P | R | F1 |
| --- | --- | --- | --- |
| SVM | 26.0 | 27.0 | 24.0 |
| Random Forest | 45.0 | 44.0 | 44.0 |
| ATLOP | 56.1 | 38.3 | 44.4 |
| DREEAM | 57.8 | 42.3 | 48.2 |
| BERT | 66.0 | 66.9 | 66.0 |
| BioBERT | 68.1 | 68.5 | 67.8 |
| SciBERT | 69.7 | 69.3 | 69.0 |
| PubMedBERT | **71.9** | **71.7** | **71.3** |
| BioLinkBERT | 71.2 | 71.2 | 70.8 |
| BioBART | 66.0 | 64.0 | 64.0 |
| Flan T5 | 45.0 | 45.0 | 40.0 |
| Llama2 13B<sup>†</sup> | 39.2 | 15.2 | 14.3 |
| Llama3 8B<sup>†</sup> | 11.3 | 11.2 | 8.34 |
| Llama 3.2-3B-Instruct<sup>†</sup> | 4.9 | 7.47 | 3.78 |
| Mistral 7B-Instruct<sup>†</sup> | 26.9 | 10.4 | 4.95 |
| BioMistral 7B<sup>†</sup> | 9.41 | 9.87 | 7.07 |

#### Simple Statistical Models: SVM and Random Forests

SVM and Random Forest exhibited notably lower performance compared to neural network models, with F1-scores of 24% and 44%, respectively. These models rely heavily on manual feature engineering and struggle to capture the complex dependencies present in document-level relation extraction tasks, particularly in the biomedical domain, where relations are context-dependent and require nuanced understanding.

#### Document-Level Models: ATLOP and DREEAM

In contrast, transformer-based models show far superior performance. Among the transformer-based models, ATLOP and DREEAM, both designed for document-level relation extraction, offer moderate results. While these models can handle overlapping relations and long-context dependencies, the lack of domain-specific pre-training on a complex task seems to penalize their performances. Moreover, ATLOP’s localized context pooling is highly sensitive to the diversity of entities within each document, and this diversity may be lacking in our use case for specific entity types.

ATLOP and DREEAM have been trained on both biomedical and general corpora, but their multi-class training was conducted using a general domain dataset. As a result, these models struggle to accurately capture the complexity of multi-class relations in the biomedical field, likely explaining their lower results.

#### Encoder-Only Models: BERT models

The nature of the dataset being dedicated to the microbiome study underlines the importance of the domain-specific pre-training for RE task. While BERT, a general pre-trained model known for its great results on such tasks, performed relatively well, it still falls behind domain-specific models like PubMedBERT and BioLinkBERT. This underscores that even highly successful general models, such as BERT, may struggle with specialized terminologies.

Transformer-based models, on the other hand, demonstrate significantly higher performance, with PubMedBERT achieving the highest F1-score of 71.3%. This is attributed to the advantages of domain-specific pre-training, as models like PubMedBERT and BioLinkBERT are trained on large-scale biomedical corpora. The attention mechanisms allows the modelling of long-range dependencies and complex interactions between entities, which are critical for relation extraction in this domain. The encoder-decoder models struggle in this domain as it leaves room for the generation of out-of-list relations.

#### Encoder-Decoder Models: BioBART and Flan-T5

Among encoder-decoder models, BioBART-base substantially outperforms Flan-T5-small, highlighting the importance of both model capacity and domain-specific pretraining for biomedical relation extraction. BioBART, pretrained on large-scale biomedical corpora such as PubMed and PMC, achieves a F1-score of 0.64. The model performs well on high-frequency relations, including part of, start, causes, and Location of. It also generalizes reasonably to mid-frequency classes such as affects, experiences, and increase. Performance degrades on low-resource or semantically diffuse relations like possible, treats, and worsen, where the model struggles to disambiguate fine-grained labels, by labeling them using broader classes. Notably, BioBART correctly predicts a few rare but well-defined relations (e.g., Negative correlation, presence), although high scores in these cases may be unreliable due to limited support. In contrast, Flan-T5-small predominantly generates the None class and rarely output general labels such as Associated with, reflecting limited generalization. These results underscore the benefit of domain-adaptive pretraining and sufficient model capacity for capturing the diversity of biomedical relations, a conclusion that is maintained withe encoder-decoder models.

#### Decoder-Only Models or Generative Models: Llama and Mistral Models

Llama2 and Llama3 showed significantly lower performance, not necessarily due to their pretraining on general domain data, but rather because of the difficulty of the task when modeled in this particular way. Both Llama versions showed the lowest performance across all metrics. This under-performance can be explained by the following elements. In the case of Llama2, the model frequently produced hallucinations, often generating relation classes that are not part of the provided list. While some of these out-of-list classes may be semantically close to the expected relations (e.g., generating “Reduce” instead of “Decrease” or “Aggravates” instead of “Worsen“), others reflect entirely different relationships that are more context-dependent (e.g., “Communication via the vagus nerve” or “Neuroprotective Effects“). This suggests that Llama2 struggles to stay within the defined set of relation classes, reflecting the issue of respecting the instructions. Additionally, Llama2 has difficulty adhering to specific output formatting rules, often producing text in a variety of random formats, further complicating the extraction process. The model also tended to focus on more general relations, neglecting the full range of more specific relationships provided, which limits its accuracy in the task that requires precise relation classification. Almost all true positives belong to the majority classes that are the most general ones.

On the other hand, Llama3, while also under-performing compared to domain-specific models, showed clearer responses, making post-processing easier and more effective. The model appears to have a better understanding of the task, with fewer hallucinations, and when hallucinations do occur, they are more closely related to one of the predefined classes on a semantic level. Moreover, Llama3 uses all the provided relation classes in its responses, indicating a better alignment with the task. However, its precision comes at the cost of generalization ability, as the model tends to predict more specific relations and struggles with broader, more general relation types, which are more prevalent in the dataset. However, it is to note that the performance decreases with the size of the models, which is underlined with Llama3.2 3B.

Similarly, Mistral 7B-Instruct and BioMistral 7B also struggled in the zero-shot setting, for different reasons. Mistral 7B-Instruct shows a strong bias toward the majority class (associated with), predicting it 284 times and thereby achieving very low label diversity and an F1 score of only 4.95%. BioMistral 7B benefits from domain-adaptive pretraining and generates a more varied set of outputs, favoring the relation increase (240 instances). However, it only predicts six out of the 23 possible relation types, reflecting limited coverage of the relations’ scope. These results further underscore the challenge of using large generative models in zero-shot biomedical relation extraction, failing to match the performance of fine-tuned discriminative approaches, particularly when precision and label granularity are essential. Further efforts should be made to adapt direct generative relation extraction to such specialized domains, for instance, by leveraging summarization-based formulations and instruction tuning, as explored in our recent work [25], where these strategies led to improved performance of generative models in the microbiome context.

Despite the strong pre-training of biomedical models, the necessity for fine-tuning remains evident, even when the fine-tuning dataset is relatively small, which is the case with MicrobioRel. It is primordial for the adaption to the sub-domain of microbiome research, and to the unique schema of relations used to annotate highly specialized datasets.

### MicrobioRel-pred: a Larger Corpus with Relation Predictions

To illustrate the relevance and utility of MicrobioRel, we report here the predictions generated by the best model of our study, the fine-tuned PubMedBERT, on a newly selected subset of 52 biomedical articles from PubTator. These articles were selected manually, from the set of articles picked using criteria as previously described in section 2, but were not pre-annotated with relations. In addition, we conducted an exploratory analysis by generating predictions for relation extraction and manually investigating a subset of these predictions related to Inflammatory Bowel Diseases (IBD) for further insights.

#### Corpus for Predictions

A set of 52 scientific articles were manually chosen based on two main criteria. We first ranked the set of articles by the average of the number of annotations of species and diseases, and subsequently selected articles and reviews published in general audience high-impact journals (e.g., Nature, Science, Cell, Cell Host Microbe) and specific aspects within gut microbiome research (i.e., host-microbe interactions, gastroenterology, cardio-metabolic diseases). Then, these 52 articles were imported from PubTator along with their annotated entities, but without the pre-defined links for relations that needed classification. To address this gap, we examined the distance between entities with established relations in the MicrobioRel-curand applied the same distribution to infer relationships within the new articles. Due to the exponential increase in possible connections across the 52 articles, we limited the number of generated links by randomly selecting a subset of relations for each document. Table 5 reports the count of overall mentions as well as the unique ones of the corpus dedicated for predictions.

**Table 5.** Named entity count on the corpus of predictions.

| Entity type | Mentions | Unique mentions |
| --- | --- | --- |
| Disease | 13 904 | 1 597 |
| Species | 19 387 | 842 |
| Chemical | 13 709 | 1 608 |
| Cell line | 206 | 43 |
| Gene | 7 942 | 784 |
| Mutation | 268 | 78 |
| **Total** | **55 416** | **4 952** |

The distribution of named entity types showed a high degree of repetition for certain entity types (such as Species and Disease), reflecting their centrality to the biomedical focus of the articles. The repetition of entities is valuable in our case, since it serves the purpose of generating a global relations graph summarizing the extracted knowledge. It provides multiple instances of relationships between key entities, helping to reinforce and validate the network of predicted interactions. Next, non-essential repeated terms were filtered out while preserving the links they form.

#### Statistical Overview of Predictions and Relation Graph

Next, an entity relation graph was generated, in which nodes represent entities and edges represent predicted relations between them, excluding the majority class *None*. This graph integrates all predicted relations within the 52 full-text articles. In addition, we defined a list of contextually descriptive yet non-essential terms, such as *human, patient*, and *mice*, which are similar to stop words in this case. The list of these terms is provided in Supplementary data. After removing these terms, we created a filtered relations graph that retained meaningful links while eliminating redundancy. With 2 987 nodes and 4 194 edges, the resulting graph shows a relatively low average node degree of 2.81, indicating that most entities have only few connections, forming loosely connected subgroups. Key entities include inflammation, dysbiosis, obesity, insulin, and butyrate, with graph node degrees of 110, 76, 74, 69, and 67, respectively. They are identified as central hubs, reflecting their frequent involvement in biomedical interactions and their relevance in microbiome research on topics such as metabolic health. The 520 connected components of the graph showed a considerable fragmentation, with many isolated subgraphs. However, the largest component, containing 2 226 nodes, represented a core structure of interconnected entities. The low global clustering coefficient of 0.0388 further suggests that the graph lacks cohesive communities, meaning that while there are important connections, few entities form closed triads.

#### Use-Case Study on Inflammatory Bowel Diseases

Inflammatory Bowel Diseases (IBD) encompasses primarily Crohn’s disease (CD) and ulcerative colitis (UC), both of which are chronic inflammatory conditions affecting the gastrointestinal tract. These diseases are characterized by symptoms such as chronic diarrhea, abdominal pain, and weight loss, and they can significantly impact the quality of life and reproductive health. Understanding the mechanisms at the onset of the disease as well as contributing to the evolution of IBD is crucial for effective management and treatment [24]. Recent research has underlined the bidirectional interactions between disease progression and changes in microbiota composition and function, pointing out the importance to study host-microbe-disease interactions [43, 16]. While gut microbiome dysbiosis and associated metabolite profiles have been clearly linked to the progression of IBD, inconsistent findings across studies are limiting our comprehensive understanding of their roles in IBD [37].

The large-scale integration of microbiome sequencing data can enable the robust identification of associations between microbes and diseases [50]. This likely holds also true for the large-scale extraction of relations and new knowledge from millions of open full-text biomedical articles. Toward this goal, recent initiatives have actually focused on the construction of an annotated corpus for microbiome knowledge base construction (MiDRED), as well as large-scale literature mining for identifying species co-occurrences (BiotXplorer) [39]. Here, mining the entity relation graph from the corpus of only 52 full-text articles, we identified key relations linking IBD, UC and CD disease entities to other entities as predicted in MicrobioRel. The resulting oriented graph of predicted relations around IBD, UC and CD (Figure 3) highlights detected relations with specific and common species and/or biomarkers associated with these conditions. While the entities IBD, UC and CD are not directly linked within this graph, IBD and CD are connected to the species Saccharomyces, which is actually a bioindicator to differentiate UC and CD, of which a specific probiotic species, *Saccharomyces boulardii*, can reduce IBD [10, 21]. Overall, this relation graph offers a first integrated overview of known associations in IBD for a very limited selection of the literature. Nevertheless, it highlights key relations between specific Enterobacteriaceae, and specifically *Escherichia coli*, and their role in gut dysbiosis associated with IBD pathogenesis and progression [1]. On the other hand, it also highlights the role of *Faecalibacterium prausnitzii* in improving IBD, and *Streptococcus* in decreasing CD, as well as the role of *Escherichia coli Nissle* in improving UC [40]. While these predictions are limited to a very small corpus of 52 full-text articles, it demonstrates the immense potential of NLP-based knowledge extraction and synthesis for enabling hypothesis generation and new discovery in microbiome research.

**Fig. 3.**
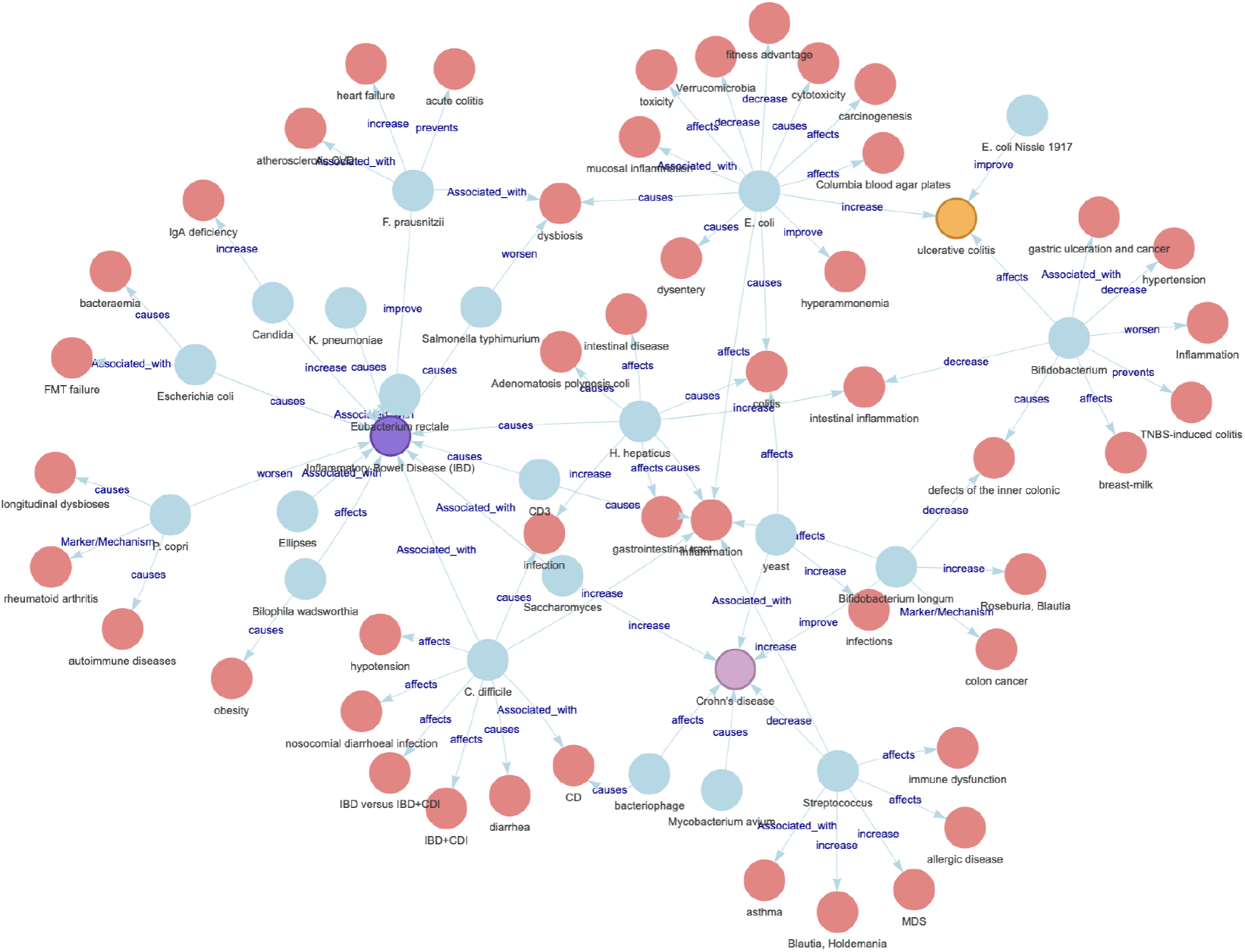
Oriented graph of predicted relations between Inflammatory Bowel Disease (purple node), Ulcerative Colitis (yellow node), and Crohn’s Disease (pink node), with associated entities on two levels, categorised by type and relation. Blue nodes represent Species and red nodes represent Diseases

## Conclusion

MicrobioRel is a novel dataset designed for document-level relation extraction in the biomedical domain. Focusing on the human intestinal microbiome, we carefully selected journal articles based on Mesh terms from a collection of PMC articles. PubTator was employed for annotating named entities related to biomedical concepts of interest. Annotation guidelines were established to refine these entities, when necessary, and to add relation annotations in the corpus. We also constructed a decision tree to simplify the labeling process, as annotating biomedical data can be a complex task. To evaluate the proposed corpus, we conducted a comparative analysis using ML-based, BERT-based, and generative models. We assessed their performance, particularly in handling imbalanced relation classes and limited size data. The approach demonstrated its utility in extracting and organizing relational knowledge from large corpora of scientific articles. By generating a knowledge graph of predicted relations, we highlight the transformation of complex biomedical textual data into a searchable and interpretable structure, facilitating further research and discovery in the field.

## Supporting information

Supplemental Experiment

## Competing interests

No competing interest is declared.

## Author contributions statement

O.E.K, S.Q. and S.C. designed the study and conceived the experiments, S.Q. and S.C. acquired funding and supervised the study. O.E.K and D.B. conducted the experiments, O.E.K, S.Q., D.B. and S.C. analysed the results. O.E.K, S.Q. and S.C. wrote and reviewed the manuscript with input from all authors.

## Acknowledgments

This work was financially supported by the ANR AIBy4 project (ANR-20-THIA-0011) and Nantes Université and performed using HPC resources from GENCI-IDRIS (Grant 2025-[AD011016540]).

## Footnotes

1 https://github.com/Stan8/MicrobioRel

2 https://www.ncbi.nlm.nih.gov/pmc/

## Notes

https://github.com/Stan8/MicrobioRel

### Competing Interest Statement

The authors have declared no competing interest.

## References

1. Valerio Baldelli, Franco Scaldaferri, Lorenza Putignani, and Federica Del Chierico. The role of enterobacteriaceae in gut microbiota dysbiosis in inflammatory bowel diseases. Microorganisms, 9(4):697, March 2021.

2. Iz Beltagy, Kyle Lo, and Arman Cohan. SciBERT:A Pretrained Language Model for Scientific Text. In Proceedings of the 2019 Conference on Empirical Methods in Natural Language Processing and the 9th International Joint Conference on Natural Language Processing (EMNLP-IJCNLP), pages 3615–3620, Hong Kong, China, 2019.

3. Olivier Bodenreider. The unified medical language system (umls): integrating biomedical terminology. Nucleic acids research, 32(Suppl 1):D267–D270, 2004.

4. Leo Breiman. Random forests. Machine learning, 45:5–32, 2001.

5. Razvan Bunescu, Ruifang Ge, Rohit J Kate, Edward M Marcotte, Raymond J Mooney, Arun K Ramani, and Yuk Wah Wong. Comparative experiments on learning information extractors for proteins and their interactions. Artificial intelligence in medicine, 33(2):139–155, 2005.

6. Hyung Won Chung, Le Hou, Shayne Longpre, Barret Zoph, Yi Tay, William Fedus, Eric Li, Xuezhi Wang, Mostafa Dehghani, Siddhartha Brahma, et al. Scaling instruction-finetuned language models. 2210.11416, 2022. arXiv preprint 2210.11416, 2022.

7. K Bretonnel Cohen, Lynne Fox, Philip Ogren, and Lawrence Hunter. Corpus design for biomedical natural language processing. In Proceedings of the ACL-ISMB workshop on linking biological literature, ontologies and databases: mining biological semantics, pages 38–45, 2005.

8. Donald C Comeau, Chih-Hsuan Wei, Rezarta Islamaj Doğan, and Zhiyong Lu. PMC Text Mining Subset in BioC: About Three Million Full-Text Articles and Growing. BMC Bioinformatics, 35(18):3533–3535, 2019.

9. Corinna Cortes. Support-vector networks. Machine Learning, 1995.

10. Guillaume Dalmasso, Françoise Cottrez, Véronique Imbert, Patricia Lagadec, Jean-François Peyron, Patrick Rampal, Dorota Czerucka, Hervé Groux Arnaud Foussat, and Valerie Brun. Saccharomyces boulardii inhibits inflammatory bowel disease by trapping T cells in mesenteric lymph nodes. Gastroenterology, 131(6):1812– 1825, December 2006.

11. Louise Deleger, Qi Li, Todd Lingren, Megan Kaiser, Katalin Molnar, Laura Stoutenborough, Michal Kouril, Keith Marsolo, Imre Solti, et al. Building gold standard corpora for medical natural language processing tasks. In AMIA Annual Symposium Proceedings, volume 2012, page 144. American Medical Informatics Association, 2012.

12. Jacob Devlin, Ming-Wei Chang, Kenton Lee, and Kristina Toutanova. BERT: Pre-Training of Deep Bidirectional Transformers for Language Understanding. In Jill Burstein, Christy Doran, and Thamar Solorio, editors, Proceedings of the Conference of the North American Chapter of the Association for Computational Linguistics: Human Language Technologies (NAACL-HLT), pages 4171–4186, Minneapolis, MN, USA, 2019.

13. Abhimanyu Dubey, Abhinav Jauhri, Abhinav Pandey, Abhishek Kadian, Ahmad Al-Dahle, Aiesha Letman, Akhil Mathur, Alan Schelten, Amy Yang, Angela Fan, et al. The llama 3 herd of models. arXiv preprint 2407.21783, 2024.

14. Oumaima El Khettari, Solen Quiniou, and Samuel Chaffron. Building a corpus for biomedical relation extraction of species mentions. In The 22nd Workshop on Biomedical Natural Language Processing and BioNLP Shared Tasks, pages 248–254, Toronto, Canada, July 2023. Association for Computational Linguistics.

15. John Giorgi, Gary Bader, and Bo Wang. A sequence-to-sequence approach for document-level relation extraction. In Proceedings of the 21st Workshop on Biomedical Language Processing, pages 10–25, Dublin, Ireland, May 2022. Association for Computational Linguistics.

16. Kerri L Glassner, Bincy P Abraham, and Eamonn MM Quigley. The microbiome and inflammatory bowel disease. J. Allergy Clin. Immunol., 145(1):16–27, January 2020.

17. Jinghang Gu, Fuqing Sun, Longhua Qian, and Guodong Zhou. Chemical-induced disease relation extraction via convolutional neural network. Database, 2017:bax024, 2017.

18. Yu Gu, Robert Tinn, Hao Cheng, Michael Lucas, Naoto Usuyama, Xiaodong Liu, Tristan Naumann, Jianfeng Gao, and Hoifung Poon. Domain-specific language model pretraining for biomedical natural language processing. ACM Transactions on Computing for Healthcare (HEALTH), 3(1):1–23, 2021.

19. Will Hamilton, Zhitao Ying, and Jure Leskovec. Inductive representation learning on large graphs. Advances in neural information processing systems, 30, 2017.

20. Kai He, Rui Mao, Qika Lin, Yucheng Ruan, Xiang Lan, Mengling Feng, and Erik Cambria. A survey of large language models for healthcare: from data, technology, and applications to accountability and ethics. arXiv preprint 2310.05694, 2023.

21. Mairead K. Heavey, Anthony Hazelton, Yuyan Wang, Mitzy Garner, Aaron C. Anselmo, Janelle C. Arthur, and Juliane Nguyen. Targeted delivery of the probiotic Saccharomyces boulardii to the extracellular matrix enhances gut residence time and recovery in murine colitis. Nature Communications, 15(1):1–16, December 2024.

22. Marĺa Herrero-Zazo, Isabel Segura-Bedmar, Paloma Martĺnez, and Thierry Declerck. The ddi corpus: An annotated corpus with pharmacological substances and drug–drug interactions. Journal of biomedical informatics, 46(5):914–920, 2013.

23. William Hogan, Andrew Bartko, Jingbo Shang, and Chunnan Hsu. Midred: An annotated corpus for microbiome knowledge base construction. In Proceedings of the 23rd Workshop on Biomedical Natural Language Processing, pages 398–408, 2024.

24. Arthur Kaser, Sebastian Zeissig, and Richard S Blumberg. Inflammatory bowel disease. Annu. Rev. Immunol., 28(1):573–621, 2010.

25. Oumaima El Khettari, Solen Quiniou, and Samuel Chaffron. Summarization for generative relation extraction in the microbiome domain. arXiv preprint 2506.08647, 2025.

26. Halil Kilicoglu, Graciela Rosemblat, Marcelo Fiszman, and Thomas C Rindflesch. Constructing a semantic predication gold standard from the biomedical literature. BMC bioinformatics, 12(1):1–17, 2011.

27. James M Kinross, Ara W Darzi, and Jeremy K Nicholson. Gut microbiome-host interactions in health and disease. Genome medicine, 3:1–12, 2011.

28. Martin Krallinger, Obdulia Rabal, Saber A Akhondi, Martın Pérez Pérez, Jesús Santamarĺa, Gael Pérez Rodrĺguez, Georgios Tsatsaronis, Ander Intxaurrondo, José Antonio López, Umesh Nandal, et al. Overview of the biocreative vi chemical-protein interaction track. In Proceedings of the sixth BioCreative challenge evaluation workshop, pages 141–146, 2017.

29. Jinhyuk Lee, Wonjin Yoon, Sungdong Kim, Donghyeon Kim, Sunkyu Kim, Chan Ho So, and Jaewoo Kang. Biobert: a pre-trained biomedical language representation model for biomedical text mining. Bioinformatics, 36(4):1234–1240, 2020.

30. Jiao Li, Yueping Sun, Robin J. Johnson, Daniela Sciaky, Chih-Hsuan Wei, Robert Leaman, Allan Peter Davis, Carolyn J. Mattingly, Thomas C. Wiegers, and Zhiyong Lu. BioCreative V CDR task corpus: a resource for chemical disease relation extraction. Database, 2016, 05 2016.

31. Carolyn E Lipscomb. Medical subject headings (mesh). Bulletin of the Medical Library Association, 88(3):265, 2000.

32. Ling Luo, Po-Ting Lai, Chih-Hsuan Wei, Cecilia N Arighi, and Zhiyong Lu. Biored: a rich biomedical relation extraction dataset. Briefings in Bioinformatics, 23(5), 2022.

33. Youmi Ma, An Wang, and Naoaki Okazaki. Dreeam: Guiding attention with evidence for improving document-level relation extraction. arXiv preprint 2302.08675, 2023.

34. Malgorzata Anna Marc, Rafal Jastrzab, and Jennifer Mytych. Does the gut microbial metabolome really matter? the connection between GUT metabolome and neurological disorders. Nutrients, 14(19):3967, September 2022.

35. Mike Mintz, Steven Bills, Rion Snow, and Dan Jurafsky. Distant Supervision for Relation Extraction Without Labeled Data. In Proceedings of the Joint Conference of the 47th Annual Meeting of the ACL and the 4th International Joint Conference on Natural Language Processing of the AFNLP, pages 1003–1011, 2009.

36. Swaroop Mishra, Daniel Khashabi, Chitta Baral, Yejin Choi, and Hannaneh Hajishirzi. Reframing instructional prompts to GPTk‘s language. In Smaranda Muresan, Preslav Nakov, and Aline Villavicencio, editors, Findings of the Association for Computational Linguistics: ACL 2022, pages 589–612, Dublin, Ireland, May 2022. Association for Computational Linguistics.

37. Lijun Ning, Yi-Lu Zhou, Han Sun, Youwei Zhang, Chaoqin Shen, Zhenhua Wang, Baoqin Xuan, Ying Zhao, Ma Yanru, Yuqing Yan, Tianying Tong, Xiaowen Huang, Muni Hu, Xiaoqiang Zhu, Jinmei Ding, Yue Zhang, Zhe Cui, Jing-Yuan Fang, Haoyan Chen, and Jie Hong. Microbiome and metabolome features in inflammatory bowel disease via multi-omics integration analyses across cohorts. Nature Communications, 14, 11 2023.

38. Claudia A Perry. Knowledge bases in medicine: a review. Bulletin of the Medical Library Association, 78(3):271, 1990.

39. Patrick Ruch, Emilie Pasche, Pierre-André Michel and Déborah Caucheteur. Biotxplorer: Navigating evidence-based biotic interactions. Biodiversity Information Science and Standards, 8:e135453, 2024.

40. Franco Scaldaferri, Viviana Gerardi, Francesca Mangiola, Loris Riccardo Lopetuso, Marco Pizzoferrato, Valentina Petito, Alfredo Papa, Jovana Stojanovic, Andrea Poscia, Giovanni Cammarota, and Antonio Gasbarrini. Role and mechanisms of action of escherichia coli nissle 1917 in the maintenance of remission in ulcerative colitis patients: An update. World J. Gastroenterol., 22(24):5505–5511, June 2016.

41. Melanie Sclar, Yejin Choi, Yulia Tsvetkov, and Alane Suhr. Quantifying language models’ sensitivity to spurious features in prompt design or: How i learned to start worrying about prompt formatting. arXiv preprint 2310.11324, 2023.

42. Isabel Segura-Bedmar, Paloma Martĺnez, and Marĺa Herrero-Zazo. SemEval-2013 task 9 : Extraction of drug-drug interactions from biomedical texts (DDIExtraction 2013). In Second Joint Conference on Lexical and Computational Semantics (^*^SEM), Volume 2: Proceedings of the Seventh International Workshop on Semantic Evaluation (SemEval 2013), pages 341–350, Atlanta, Georgia, USA, June 2013. Association for Computational Linguistics.

43. Yue Shan, Mirae Lee, and Eugene B Chang. The gut microbiome and inflammatory bowel diseases. Annu. Rev. Med., 73(1):455–468, January 2022.

44. Maxim Tkachenko, Mikhail Malyuk, Andrey Holmanyuk, and Nikolai Liubimov. Label Studio: Data labeling software, 2020-2022. Open source software available from https://github.com/heartexlabs/label-studio.

45. Hugo Touvron, Louis Martin, Kevin Stone, Peter Albert, Amjad Almahairi, Yasmine Babaei, Nikolay Bashlykov, Soumya Batra, Prajjwal Bhargava, Shruti Bhosale, et al. Llama 2: Open foundation and fine-tuned chat models. arXiv preprint 2307.09288, 2023.

46. Vladimir Vapnik. Statistical learning theory. John Wiley & Sons google schola, 2:831–842, 1998.

47. Lu Wang, Shiqi Wang, Qing Zhang, Chengqi He, Chenying Fu, and Quan Wei. The role of the gut microbiota in health and cardiovascular diseases. Mol. Biomed., 3(1):30, October 2022.

48. Chih-Hsuan Wei, Alexis Allot, Robert Leaman, and Zhiyong Lu. Pubtator central: automated concept annotation for biomedical full text articles. Nucleic acids research, 47(W1):W587–W593, 2019.

49. Ye Wu, Ruibang Luo, Henry CM Leung, Hing-Fung Ting, and Tak-Wah Lam. Renet: A deep learning approach for extracting gene-disease associations from literature. In Research in Computational Molecular Biology: 23rd Annual International Conference, RECOMB 2019, Washington, DC, USA, May 5-8, 2019, Proceedings 23, pages 272–284. Springer, 2019.

50. Liwen Xiao, Fengyi Zhang, and Fangqing Zhao. Large-scale microbiome data integration enables robust biomarker identification. Nat. Comput. Sci., 2(5):307–316, May 2022.

51. Zhaohui Yan, Zixia Jia, and Kewei Tu. An empirical study of pipeline vs. joint approaches to entity and relation extraction. In Yulan He, Heng Ji, Sujian Li, Yang Liu, and Chua-Hui Chang, editors, Proceedings of the 2nd Conference of the Asia-Pacific Chapter of the Association for Computational Linguistics and the 12th International Joint Conference on Natural Language Processing (Volume 2: Short Papers), pages 437–443, Online only, November 2022. Association for Computational Linguistics.

52. Michihiro Yasunaga, Jure Leskovec, and Percy Liang. LinkBERT: Pretraining language models with document links. In Smaranda Muresan, Preslav Nakov, and Aline Villavicencio, editors, Proceedings of the 60th Annual Meeting of the Association for Computational Linguistics (Volume 1: Long Papers), pages 8003–8016, Dublin, Ireland, May 2022. Association for Computational Linguistics.

53. Hongyi Yuan, Zheng Yuan, Ruyi Gan, Jiaxing Zhang, Yutao Xie, and Sheng Yu. BioBART: Pretraining and evaluation of a biomedical generative language model. In Proceedings of the 21st Workshop on Biomedical Language Processing, pages 97–109, Dublin, Ireland, May 2022. Association for Computational Linguistics.

54. Dongxu Zhang, Sunil Mohan, Michaela Torkar, and Andrew McCallum. A distant supervision corpus for extracting biomedical relationships between chemicals, diseases and genes. In Proceedings of the Thirteenth Language Resources and Evaluation Conference, pages 1073–1082, Marseille, France, June 2022. European Language Resources Association.

55. Wenxuan Zhou, Kevin Huang, Tengyu Ma, and Jing Huang. Document-level relation extraction with adaptive thresholding and localized context pooling. In Proceedings of the AAAI conference on artificial intelligence, volume 35, pages 14612–14620, 2021.

