## Supplemental Experiment for "MicrobioRel: A Set of Datasets for Microbiome Relation Extraction"

### Supplementary data

#### 1 Preliminary Experiments

In this appendix, we provide an in-depth discussion of various fine-tuning frameworks that have been employed to address the challenges posed by our highly imbalanced and specialised dataset. Firstly, the imbalance problem was addressed by implementing class weights during the training of BERT-based models, with the aim of adjusting the learning process according to the frequency of each relation type. In addition, we investigate a common practice in relation extraction that involves masking the entities involved within each triplet by replacing them with their respective types. This technique is intended to refocus the model's attention from the specific terminology of the entities to the semantic relation they share.

##### 1.1 Experimental Setup

Given the nature of the dataset, which is heavily imbalanced and specialised, we discuss several fine-tuning frameworks. First, we aimed to address the imbalance issue by introducing class weights during the training of the BERT-based models. This approach was designed to adapt the learning process to the frequency of each relation type. However, while this method favoured the majority classes, it did not significantly improve the performance of the minority classes, which are crucial for highlighting the most precise relations and studying information novelty. Therefore, we decided to prioritise the results of the regular fine-tuning instead.

We also explored a common practice in relation extraction that involves masking the participating entities in each triplet (entity1, relation, entity2) by replacing them with their types. This effective technique aims to redirect the focus of the model from the specialised vocabulary of the entities to the semantic connection between them. Although we experimented with incorporating the six entity masks into the vocabulary, this approach did not improve the results. This suggests that while masking can be beneficial, its effect may vary depending on the specific context and dataset. These results are presented below. Conversely, preliminary experiments showed that masking entities in generative approaches was ineffective. Masking the entities in the prompts led to worse results than to using the original paragraphs.

#### 1.2 Assessment of Fine-Tuning Strategies for Imbalanced Biomedical Relation Extraction

For the BERT-based models, the experiments were conducted within different fine-tuning frameworks. In addition to the conventional fine-tuning of the models (denoted "Reg FT"), one approach involved incorporating the 6 entity masks into the vocabulary prior to training (denoted "AV + FT", for "Added Vocabulary + Fine-Tuning"). Finally, class weights were introduced to address the issue of imbalanced data (denoted "FT+CW", for "Fine-Tuning with class weights").

In initial experiments, we fine-tuned the cased version of the aforementioned models over 12 epochs, using a fixed batch size of 8 and a learning rate of $1 \times 10^{-5}$.

| **Models** | **Reg FT** | **AV+FT** | **FT+CW** |
|---|---|---|---|
| BERT-*base* | 42.6 | 36.8 | 22.1 |
| BioBERT | 51.4 | 44.5 | 38.4 |
| SciBERT | **54.6** | **42.6** | **46.6** |
| PubMedBERT | 48.0 | 36.2 | 25.3 |

Table 1: F1-scores (in %) on the test set, for 4 models and various fine-tuning settings.

The results in Table 1 illustrate the complexity involved in adapting relation extraction methods to a highly imbalanced and specialised dataset. For the BERT-based models, regular fine-tuning generally outperformed the class-weighted approach, particularly in capturing minority classes. Although class weights helped the model to recognise majority classes, they had minimal impact on improving the accuracy of rare relations. For example, the relation Presence, which occurs only once in the three splits, was only recognised by SciBERT using the regular fine-tuning. The inclusion of masks showed mixed results. While it occasionally helped to refine the focus on relational patterns, its application was inconsistent across different model types.

#### 2 Prompt Template

| | |
|---|---|
| **Instruction:** | Generate the class of the relation between "Parkinson's disease" starting in character 259 and "motor deficits" starting in character 345 from the Classes list based on the input. |
| **Constraints:** | You have to output the class only. Justification and explanation are prohibited. Classes: Increase, Decrease, Stop, Start, Improve, Worsen, Presence, Negative_correlation, Affects, Causes, Complicates, Experiences, Interacts_with, Location_of, Marker/Mechanism, Prevents, Reveals, Treats, Physically_related_to, Part_of, Possible, Associated_with, None. |
| **Input:** | Patients suffering from PD have different bacterial flora compared with healthy controls. Microbiota have an impact on disease progression, which has been established in the fecal-transplantation experiments. Administration of the microbiota of patients with Parkinson's disease to mice showed the development of neuroinflammatory processes and motor deficits. |
| **Output:** | - |

Table 2: Example of the zero-shot setting prompt in relation extraction, following the described prompt template.

#### 3 MicrobioRel-cur

Another aspect of relations extraction involves assessing the participating entities for each relation label. Conforming to the annotation guidelines, some of the relations only occur between specific entity types, such as the relation labels Treats, or Complicates. In contrast, broader relations occurring across all entity types are more likely to be prevalent. As shown in Figure 1, the relative contribution of a given relation is in agreement with the number of entities. This observation validates the hypothesis posited in document selection: for a relation to be conveyed in a text, it is imperative that a minimum of two named entities are referenced within the same context.

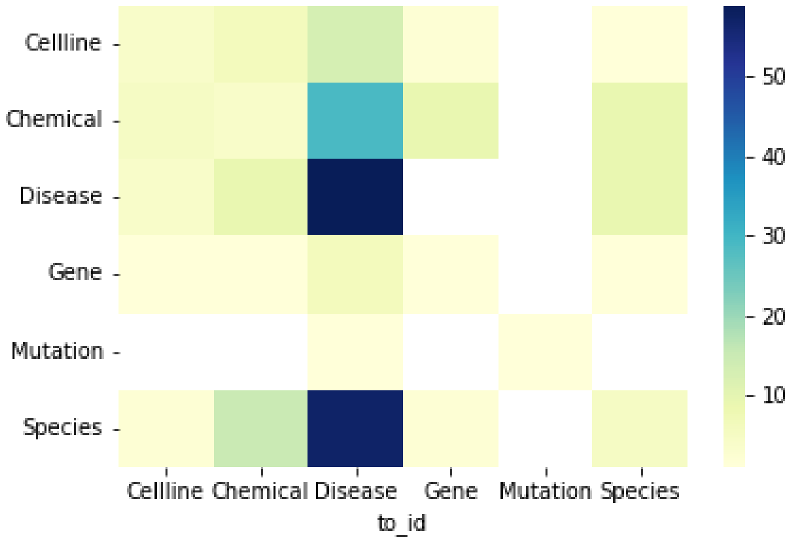


Figure 1: Heatmap representing pairs of entity types participating in relations. The color gradient indicates the number of times each entity pair appears together in a relation across the annotated corpus, with darker colors corresponding to higher frequencies.

### 4 List of Removed Entities Before Graph Creation

Prior to the construction of the relation graph, the following list of entities, highly prevalent in the MicrobioRel-predcorpus but less relevant for IBD, was generated and their occurrences were removed from the dataset:

- Human-Related Entities: *women*, *men*, *patients*, *human*, *people*, *participants*, *human microbiome*, *human gut microbiota*, *child*, *children*, *infant*, *infants*, *participant*, *man*;
- Animal-Related Entities: *mice*, *mouse*, *murine*, *rat*, *rats*, *cat*, *cats*, *rabbit*, *rabbits*, *sheep*, *sheep*, *cow*, *cows*, *Kangaroo*, *transgenic*, *drosophila*, *donor*;
- Microbiome and Metagenome-Related Entities: *gut*, *gut microbiome*, *human gut microbiota*, *gut metagenomes*, *gut metagenome*, *metagenome*, *metagenomes*;
- Miscellaneous/Generic Entities: *prions*, *prion*, *increase infection*, *shuttle vector*.

While these entities were important for providing contextual information according to the annotation guidelines, they were not necessary for understanding the interactions between the primary entities of interest. Specifically, these entities serve to introduce context, particularly in the case of the Experiences relation, but do not directly contribute to the core relations being analysed. By excluding these entities, the resulting graph of extracted relations focus on the relevant interactions and discard generic entities widely used in scientific articles, ensuring a more targeted representation of the key entity interactions.
